# Genetic diversity evaluation among fifty sugarcane accessions from the Zimbabwe Sugar Industry using DNA-SSR markers and qualitative traits

**DOI:** 10.1101/2020.01.20.909028

**Authors:** E Mugehu, L.T. Mpofu, P Manjeru

**Affiliations:** Midlands State University, P. Bag 9055, Gweru, Zimbabwe; Zimbabwe Sugar Association Experiment Station, P. Bag 7006. Chiredzi, Zimbabwe

**Keywords:** genetic diversity, morphology, SSR markers

## Abstract

Due to the increasingly diverse needs of the Zimbabwe Sugar Industry, the need for developing improved varieties of sugarcane cannot be overemphasized. Since the establishment of experiment station in 1966, the Zimbabwe Sugar Industry (ZSI) has always been dependent on importation of crosses from other countries for development and release. However, the ZSI is now planning on developing its own crossing facilities. The ZSI’s gene bank has up to 746 imported varieties which can be used as potential parents in a breeding program. With the impending establishment of Zimbabwe’s own crossing facilities, morphological and molecular characterization of the available germplasm is essential to identify parental lines which can be used during crossing. The main objective of this study was to determine the genetic and phenotypic diversity of fifty ZSAES gene bank sugarcane varieties (accessions) and assess how the existing diversity can be utilized in a breeding program in parental selection and variety identification. Fifty sugarcane accessions were chosen based on country of origin and availability of seed and evaluated for qualitative morphological traits followed by molecular evaluation using DNA-SSR markers in a PCR based protocol. Fifty ‘one eyed’ setts were planted in the greenhouse and transplanted into labeled asbestos pots with ZSAES media. Fresh leaf samples were collected from the plants for DNA extraction using a modified CTAB method. Field trials for morphological assessment were laid out in a completely randomized design in a 1.5 × 4m plot. Agglomerative, hierarchical UPGMA clustering based on morphological traits with SAS^®^ (9.3) categorized the fifty accessions into two main groups. The morphological traits managed to classify the accessions according to pedigree data, source of origin and similarities in traits like internode waxiness and adherence of leaf sheath. Contrastingly, molecular UPGMA cluster analysis by DarWin^®^ (6.0) grouped the accessions into three main clusters. The three clusters were aligned according to time of release of varieties with the latter varieties in cluster III and the older ones in cluster I. The groups also showed clustering according to similarities in pedigree and source of origin. The molecular and morphological phylogenetic clusters suggest horizontal gene transfer through parallel and convergent evolution. Similarity trends observed also highlight similarities in breeding goals in different parts of the world. The information obtained from the study justifies the need for more importation of varieties from across the world to broaden the genetic base of the germplasm pool in the ZSI’s breeding program. Information from morphological data can be used for identification of released varieties in the ZSI using unique descriptors. More varieties should be analyzed and agronomic characterization should also be done to complement the results of this study and be incorporated into a database.

## Introduction

Sugarcane has various different end products in the ZSI and the environmental conditions across the industry differ with geographical location. This calls for the release of high sugar producing varieties that also have other several desired agronomic traits and environmental adaptations. This is because the variety is the pivot around which entire production system revolves. Therefore, scientific sugarcane cultivation must involve choosing and developing an appropriate variety for the agro-climatic zone, prevalent pest, diseases, soil type and end product. Improved varieties are now available for different growing conditions in the world and new varieties are continuously developed and released by sugarcane research centers world over. The Zimbabwe Sugar Industry (ZSI) has never made its own crosses due to the lack of ideal daylight temperatures and day lengths which are conducive for flowering. Floral initiation in sugarcane is induced by a small decrease (30 to 60 s-^1^ d-^1^) in day length below 12 h 30 min (Berding 1995; Moore and Nuss 1987). Such conditions are not available in Zimbabwe and as a result, artificial photoperiod is needed to aid sugarcane flower initiation in breeding genotypes by advancing the flower initiation phase to an earlier and more conducive environment (>18.0 °C and <32.0 °C) (Nuss and Berding 1999). As a result, the ZSI has always been dependent on the importation, testing and release of varieties due to the lack of photoperiod facilities which would enable cross pollinations to be done locally.

However, the ZSI is now planning on developing its own crossing facilities. This would enable total control of the parental lines to be used in the cross and the environment for adaptation. Crossing facilities would adequately suit the industry as it is sitting on a reserve of up to 746 varieties imported from different countries which are being conserved in the Zimbabwe Sugar Association Experiment Station’s gene bank. These varieties are a rich source of germplasm which can be used as parental lines in crosses. Characteristics of all the varieties that are being commercially grown and also conserved in that industry’s gene bank would be greatly useful in parental selection. Characteristics should be identified using morphological and molecular techniques. The use of molecular markers to characterize the varieties in tandem with phenotypic characterization would enable the varieties to be classified into groups so that the breeder can efficiently select parents for crosses. Parents’ choice is the first step in plant breeding program through hybridization in sugarcane. In order to benefit transgressive segregation, a greater genetic distance between parents is necessary (Sajib *et. al*, 2012). Molecular markers would also confirm differences which may not be seen at phenotypic level. Reliance on morphological characteristics and pedigree history alone does not give a sound inference on the genetic structure and diversity of a population. Molecular characterization of the genotypes also gives precise information about the extent of genetic diversity which helps in the development of an appropriate breeding program (Sajib *et. al*, 2012). Simple sequence repeats (SSR) are ideal molecular markers for genetic variation identification of germplasm (Powell *et al*., 1996; Ma *et al*., 2011). The use of SSRs for analysis of germplasm diversity and utilization of heterosis is ultimate, especially in identification of species with closer genetic relationship such as, sugarcane (Zhou *et al.*, 2003; Jin *et al.*, 2010; Ma *et al*., 2011). The main objective of this study was to evaluate the genetic diversity of 50 sugarcane accessions using 20 SSR markers and 20 qualitative traits from the Zimbabwe Sugar Industry and various other countries so as to assess how the accessions’ diversity can be utilized in the breeding program in terms of parent line selection and variety identification.

**The specific objectives of the study were to:**

- Determine the genetic diversity among the 50 sugarcane varieties using molecular DNA-SSR markers in a PCR-based protocol.
- Phenotypically characterize the diversity present amongst 50 sugarcane accessions under field conditions based on their qualitative traits.
- Group the varieties into clusters according to similarities in their genome and morphology.

### Description of the study area

The trial was carried out during the 2016-2017 sugarcane growing season at the ZSAES in the South-Eastern lowveld of Zimbabwe. The ZSAES is located centrally in the Zimbabwe Sugar Industry being approximately 99km along the Ngundu-Tanganda road with latitude 20°01’ S and longitude 28° 38’ E. The experiment station is situated on sandy loam soils classified as lithosols 430 meters above sea level. The ZSAES cane area is mainly under both overhead and surface irrigation with an average rainfall of 625mm per annum (20-year mean) falling predominantly in the hot summer months (October-March). The winters are relatively cool and dry with mean air temperatures varying from 26° C (October-January) to 16° C (June-July).

### Germplasm selections and Experimental procedures

Fifty sugarcane varieties were chosen to represent part of the germplasm broadly cultivated in the South-eastern Lowveld of Zimbabwe and available to the alcohol and sugar factories (Table 2). Choice of varieties was based on availability of planting material and country of origin. The set included sugarcane varieties from breeding programs from various countries in the world and also released varieties from the Zimbabwe Sugar Association Experiment Station itself. The planting material was collected from the germplasm source in the Zimbabwe Sugar Association Experiment Station gene bank. All field experiments were laid out in a completely randomized design (CRD) with the varieties as treatments. Treatments were randomly allocated to plots with net plot size being 2 rows by 4meters at 1.5m interrow spacing with a two peripheral row discard on either side of the field. Planting was done on the 29^th^ of November 2016 as double planting with planting material comprising of 11 stalks per plot. A variety of common knowledge (ZN10) was included after every 9th candidate variety under study to enable easy assessment of distinctness between genotypes. Fertilizer applications were based on soil analyses results. Nitrogen was applied in three splits at four weekly intervals at the rate of 47kgN/ha, 46kg/ha and 46kg/ha respectively. Muriate of Potash was applied in two splits at four weekly intervals at 97kg/ha each split. Integrated weed management was done which included the use of pre-emergence herbicide Acetochlor (2l/ha), Metribuzin (2l/ha) and Servian (Halosulfron-methyl) at 7g/15litres of water. Insect pest control included Regent 200SC (Fipronil) for termites and Dursban 480 EC for Black maize beetle control. Scouting routines for insect pests, pathogens and diseases were done regularly and infected plants were rouged.

### Morphological data collection

Morphological traits were measured according to the UPOV (2005) international guidelines after the grand growth stage at 8 months after planting (July 2017). Six stalks were taken at random from different stools in the plot. Known varieties were used to clarify the state of expression for each characteristic. States of expression given for each characteristic were used to define the characteristic and to harmonize descriptions (Table 1). Each state of expression was given a corresponding numerical note for ease of recording of data and for the production and exchange of the description. Traits like plant growth habit, leaf blade curvature, tillering and the adherence of the leaf sheath were taken before destructive sampling of stalks was done. For the rest of the traits, stalks were taken from the field and laid out on to a large table for examination. All colors were examined visually and scored according to the Royal Horticultural Society (London) color chart (Gashaw *et al.*, 2016).

**Table 1:**
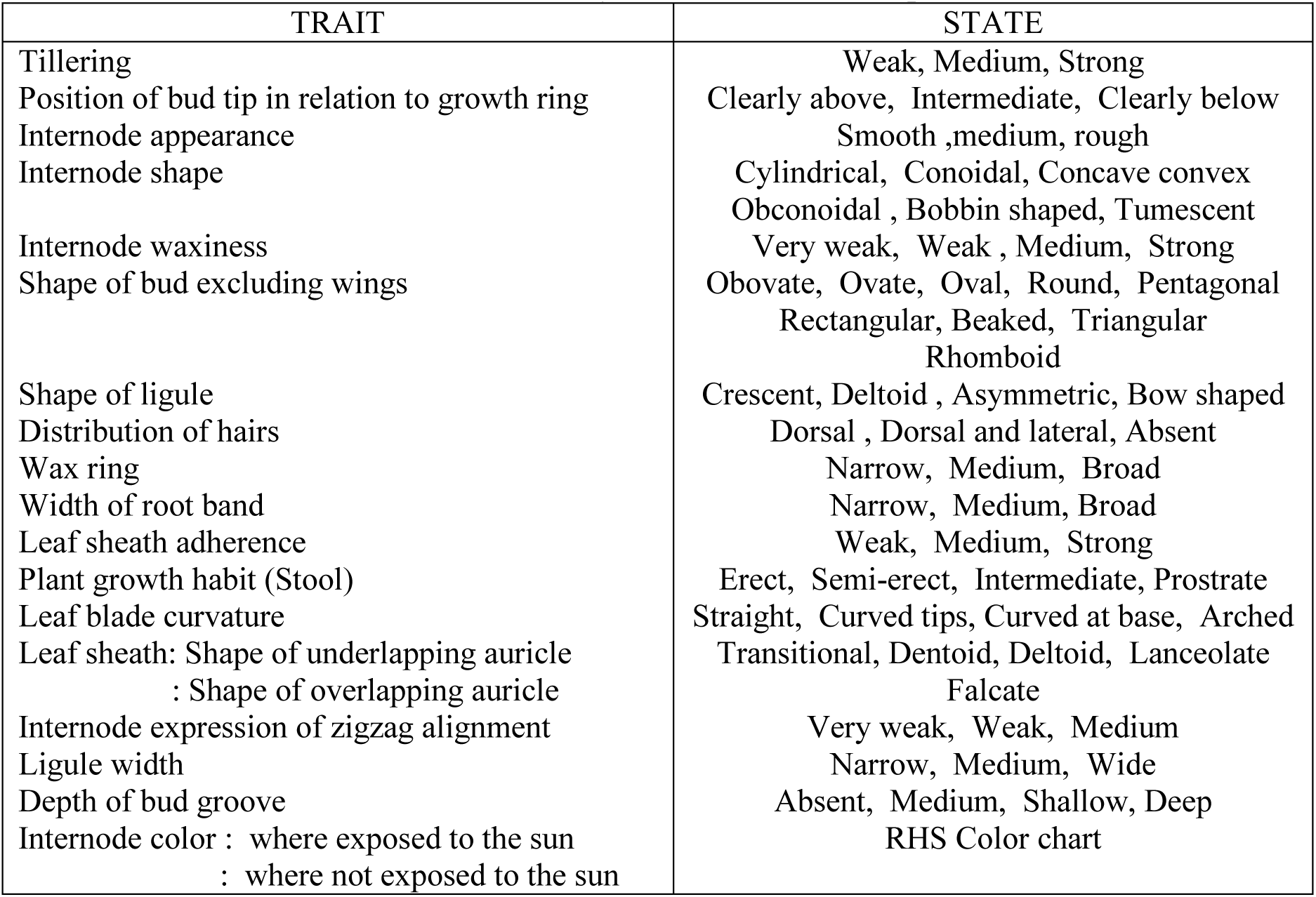
Qualitative traits used in the study and their states of expression.

**Table 2:**
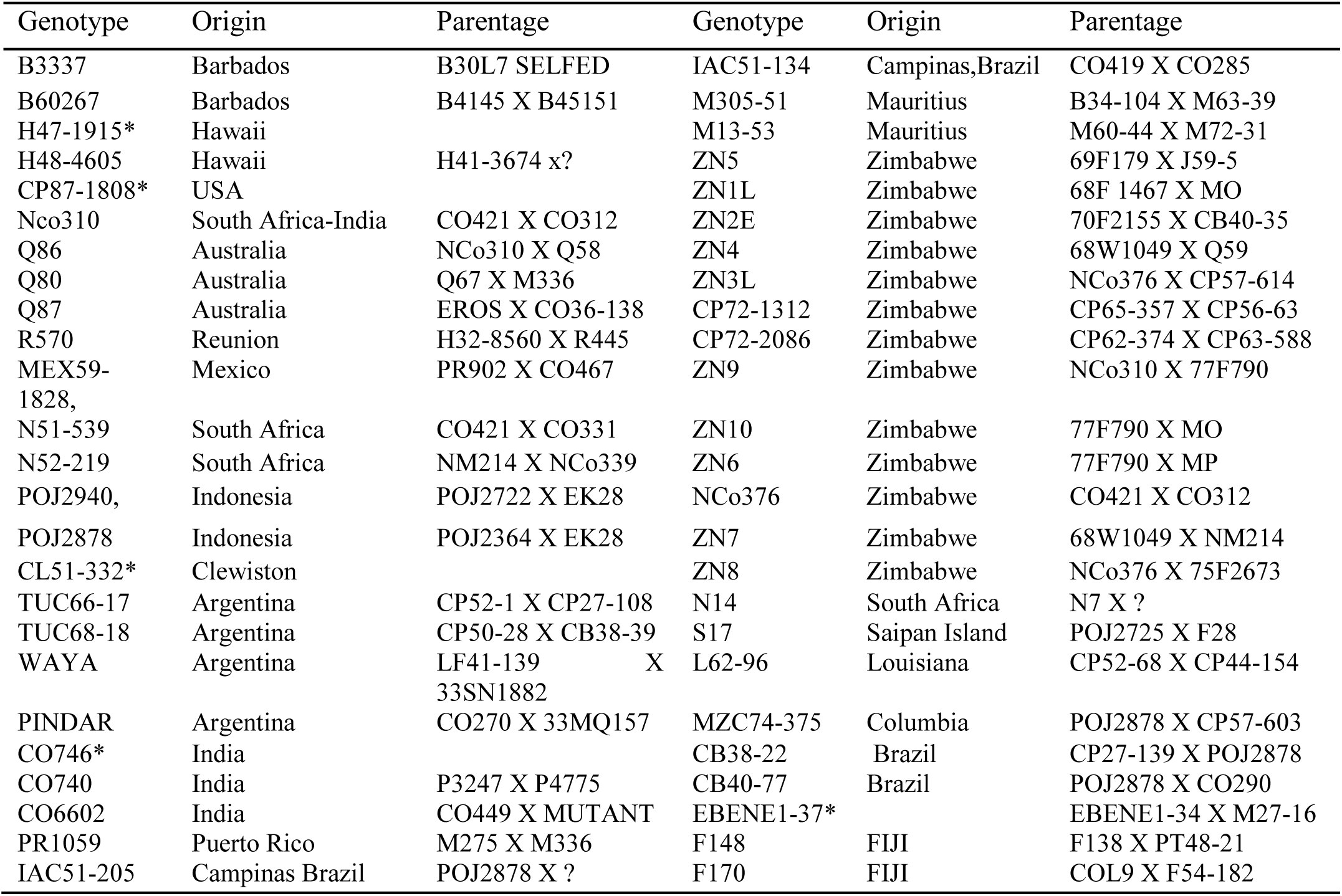
List of sugarcane varieties used in the study including source and parentage. Information on sugarcane parentage obtained from CIRAD database (*tropgenedb.cirad.fr*)

### Leaf material and DNA extraction

Fifty varieties were planted in the greenhouse. Two stalks were taken per variety and cut into one-eyed setts. Setts were dipped into a fungicide (Shavvit) at 1ml: 1litre of water for 5 minutes before planting. The setts were laid out on saturated ZSAES media which consisted of top soil, sandy soil and organic manure at the ratio of 5:2:2 respectively. The plants were fertigated with Hoagland’s nutrient solution once a week and transplanted into labeled asbestos pots containing ZSAES media. Plant DNA extraction was done using the Cetyl Trimethyl Ammonium Bromide (CTAB) method as described by Doyle and Doyle (1987) with slight modifications in relation to the use of liquid nitrogen to facilitate membrane maceration. Leaf material of approximately 0.1mg from samples of the varieties was weighed and washed with sterile (double distilled, ultraviolet light and reverse osmosis treated) water. The leaves were submerged and fixed in 95% ethanol alcohol for 1 hour in a sterile petri dish following the protocols by Sharma *et al*., (2010) and Biswas *et al*., (2011) instead of grinding them in liquid nitrogen. This was prompted by the unavailability of liquid nitrogen locally and also its need for careful handling to avoid vaporization. After submersion in ethanol, the leaves were transferred to a sterilized, pre-chilled mortar and ground to a fine powder using a ceramic pestle. The homogenized powder was then transferred to a 1.5 ml polypropylene centrifuge tube containing 1.0 ml of pre warmed (62°C) 2X CTAB buffer. The samples were then subjected to a series of incubation, centrifugation steps as described by Doyle and Doyle (1987).

### DNA Quantification

Suspended DNA was first checked for integrity and quality using agarose gel electrophoresis. One gram of agarose was dissolved in 100ml of 1 × Tris-Borate-EDTA (TBE) buffer by heating with gentle swirling to form a semi-solid gel. The gel slab was then placed in an electrophoresis tank containing 700ml of 1×TBE buffer and 35µL of ethidium bromide. 8uL of the extracted DNA was mixed with 2µL of 6x gel loading dye (0.25 % (w/v) bromophenol blue, 0.25 %(w/v) xylene cyanol, 30% (v/v) glycerol) on sterile laboratory film and loaded into the gel wells using a micropipette. The gel was then run on an electric current of 100 Volts for 45 minutes on a Cleaver Scientific® 300V horizontal gel electrophoresis system. Genomic DNA was then visualized on a Cleaver Scientific® Clearview UV transilluminator to determine the integrity and purity of the DNA. The DNA quality was assessed by measuring the absorbance of a 1:10 dilution of the sample. 1ml of Tris EDTA (TE) buffer was transferred into a cuvette and read onto a Libra Biochrom® S80PC UV/VIS at 260nm, 280nm and 320nm wavelengths 5µL of the genomic DNA was then pipetted into 995uL of TE buffer and absorbance readings were also taken at 260nm, 280nm and 320nm for each of the tubes. DNA purity was checked using the ratio A260/A280 to give an estimate of protein contamination. If the ratio was close to 1.8, the DNA was considered to be relatively pure.

### Polymerase Chain Reaction (PCR)

PCR reactions were run on Cleaver Scientific® Multigene Optimax thermocycler. PCR optimization and conditions was referenced both from Pan *et al* (2000) and through gradient PCR using primer sequences agreed upon by the sugarcane microsatellite consortium (Cordeiro *et al*, 2003). Each 10µL reaction contained 25 ng genomic DNA; 5.0 μL 10X reaction buffer (20 mM Tris-HCl, pH 8.0, 0.1 mM ethylenediaminetetraacetic acid (EDTA), 1.0 mM dithiothreitol, and 50% (v/v) glycerol); 25 mM MgCl_2_^+^; 0.1 mM each dATP, dGTP, dCTP, and dTTP; 0.5 μL each primer (Forward and Reverse primers); and 1 unit Taq polymerase (New England Biolabs). The PCR conditions were as follows: initial denaturation at 95°C for 15 minutes; 40 cycles of 15 seconds at 94°C, 15 seconds of each primer’s specific melting temperature and 1 minute of extension at 72°C. The final extension step was 10 min at 72°C and a reaction hold at 4°C.

### Gel Electrophoresis

A total of 8μL of each sample was separated by electrophoresis on a 2% agarose gel (50% agarose and 50% agarose Metaphor, CAMBREX) containing 1X TBE buffer and 0.5 μg/mL ethidium bromide solution 90 Volts for 1 hour. A 1-kb ladder (New England Biolabs) was used as the weight molecular marker and loaded on the first and last well. A negative control was included in one well where loading dye was loaded without any amplification product to check for well contamination. Gels were then visualized under UV light using a Cleaver Scientific gel transilluminator. The image was captured using a Nikon D3200 photographing system for further allele scoring and analysis.

### Data Analysis

Allele numbers per locus from SSR analysis were determined based on their relative positions in the gel. The data was analyzed using band to band gel readings with a reproducible band signifying the presence of a particular allele and was given a score of 1, and its absence was given a score of 0 in each sample. The alleles were also scored as an integer of their base pairs. A rectangular binary data matrix was constructed and analyzed using the DARwin software version 6.0.14 (Singh *et al.*, 1999). The dissimilarity matrix was constructed using simple pairwise matching algorithm and Euclidean distances were calculated using allelic data (Sokal and Sneath, 1963). A dendrogram was constructed based on genetic distance using the Unweighted Neighbor-Joining method and bootstrap analysis. Coefficient of dissimilarity for morphological data was calculated in all of the pairwise comparisons of the sample germplasm based on squared Euclidean distances. A dendrogram was created by cluster analysis with unweighted pair grouping method based on arithmetic averages (UPGMA) using the computer program SAS^®^ 9.3 (SAS Institute, Cary NC), (Gashaw *et al*., 2016).

## Results and discussion

Morphological data grouped the fifty accessions into two main clusters and two solitary outliers (Figure 1). Cluster I consisted of 17 accessions which were bunched according to mainly pedigree(parental) history as well as the country of origin for instance varieties Q86, ZN3L and N51-539 are of Indian descent. There were also four varieties namely M13-53, EBENE1-37 and PR1059 which have parents from Canal Point, America. The three released varieties, ZN10, ZN1L and ZN5 all have Formosan parents (69F179, 68F1467 and 77F790 respectively). Additionally, R570 is a cross between a Hawaiian (H32-8560) and a French variety (R445) which perhaps is why it was inclined near H47-1915 in that cluster. MEX59-1828 is of Puerto-Rican and Indian parentage hence in the same cluster with PR1059 and other varieties like N51-539 which are also of Indian parentage. Cluster II, however, was characterized by accessions discordantly grouped without any pattern of grouping by country of origin, time of release or pedigree history. Cluster II has four varieties from India (Co), seven varieties from South Africa (N), two varieties from Clewiston, America (CL), two varieties from Brazil (IAC) and three Argentine varieties (TUC, WAYA and PINDAR). Although grouping patterns in this cluster were discordant, there appeared to be some common trends in regards to indirect parents and source of origin of the parents.

**Figure 1:**
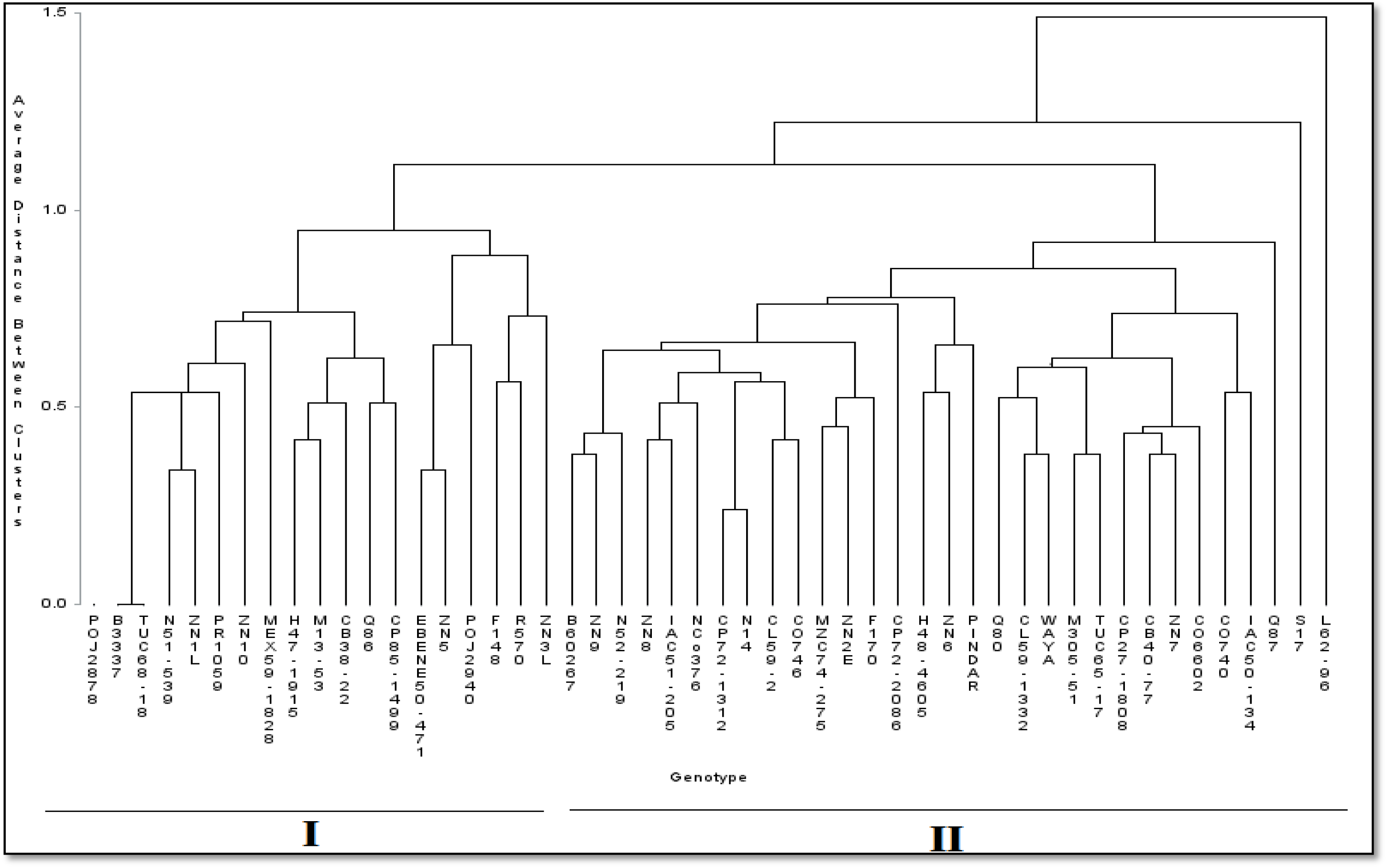
A phylogenetic tree showing clustering patterns after morphological analyses (SAS 9.3).

An outlier was observed, a variety from Louisiana, L62-96 with parents CP52-68 and CP44-154. Despite it having parents of American origin (Canal Point), L62-96 was not found in either cluster one which had a lot of Canal Point varieties or cluster II which also had some American varieties from Clewiston. Another outlier, S17, which is a product of a cross between POJ2725 and F28, was also out of articulation from the other Java (POJ) and Fiji varieties. The low morphological diversity observed among fifty accessions maybe used as an indicator of the presence of genetic uniformity of the germplasm in the gene bank. Given that from these genotypes potential parents in cross pollinations are going to be drawn, the sample size must be increased for future studies so that more groupings can be afforded. If the same pattern of uniformity is observed in a bigger sample, collecting missions must be resumed to import more diverse varieties to widen the genetic base. The similarities in morphology which were being observed may evidently be explained by the fact that genotypes have shared parentage and country of origin. Genotypes from the same country of origin would be adapted towards the same environment hence similarities would be evident in their morphology. Therefore, evolution would occur in the same direction due to the incidence of the same selection pressure. For instance, selection pressures like drought and pests incidence would call for the plant to have particular morphological and physiological characteristics to survive. Morphology is dependent on physiological processes in development and the function of the physiological characters depends on the morphology of the individual (Aljanabi *et al*., 1999). If those physiological processes are subjected to selection pressure and are forced to adapt, the morphology of the plant also changes in that direction.

Furthermore, most morphological and physiological characteristics are controlled by multigene families which are of ancient origin for instance the homeobox genes which are similar at class and order level (McCune and Grace, 2002). This suggests that the grand stages of development are controlled by the same or similar sets of genes in many different varieties hence resulting in similar morphologies. Additionally, although the breeding institutes might differ in one country, breeding goals are most likely to be similar because the biotic and abiotic conditions for the sugarcane growing areas would be similar or close. This means that two genotypes from the same country but from different breeding institutes can be morphologically similar as seen by varieties from Clewiston and Canal Point, America which are in the same cluster.

The results from the cluster analyses point towards particular patterns of evolution in sugarcane history which include convergent evolution, cladistics, parallel evolution and homeomorphy. Homeomorphy entails the occurrence of two or more genotypes that are morphologically similar although relatively ‘distantly’ related and this occurrence is due to convergent evolution (McCune and Grace, 2002). Convergent evolution is a common occurrence where genotypes which are not monophyletic (not closely related) in terms of their pedigree, evolve similar traits as a result of having to adapt to similar environments or ecological niches. This is evident in the clusters which contained some genotypes which do not share common ancestry or parentage whatsoever. For instance, the alignment of B3337 which is a product of selfing of B30L7 in cluster I. Genotype B3337 does not share direct or indirect parents with the rest of the members of the cluster but it is aligned with those members due to similarities in morphology. Discordant presence of genotype F170 in cluster II confirms cladistics and convergent evolution. Evolution could have occurred in the same direction due to reasons other than similarity in ancestry and parentage like natural selection, genetic drift and neutral mutations. Phenotypic diversity is a product of accumulation of novel mutations and their conservation that has facilitated adaptation towards different environments. These mutations could have been incorporated into the genome by natural selection. In this case, the genotypes which are showing similarities are from different parents and different sources of origin which hints on similar selection pressures in different locations. The breeder’s breeding objectives may be similar to another breeder’s who is in a completely different geographical location hence resulting in selection towards a similar crop ideotype indicating parallel evolution.

Molecular analyses produced three major clusters with 6 sub clusters (Figure 2). Varieties of the same geographical origin were falling into the same cluster and sub cluster for instance Fiji (F148 and F170), Brazil (CB30-77, CB60-27, IAC51-205), Indonesia (POJ2878, POJ2940). The twelve released varieties from Zimbabwe which are currently commercially grown also fell into the same cluster and subcluster except for ZN9. Early release varieties from the pre-1966 era including Brazilian (CB, IAC), some Indian (Co), Indonesia (POJ) varieties fell into the same cluster I which included French (R570) and ancient Argentine varieties (TUC). Cluster II mostly included varieties released after those in Cluster I consisting of Australian (Q, PINDAR), Hawaiian (H), Indian and South-African (N) varieties. American (CP) varieties were found in all three clusters of the dendrogram. All the Zimbabwe-released varieties that are currently being commercially grown in the Zimbabwe Sugar Industry were grouped together in cluster III. The released varieties were still bunched together within the sub clusters with NCo376 being flanked by ZN8 and ZN3L. N14 and ZN1L were aligned towards cluster II.

**Figure 2:**
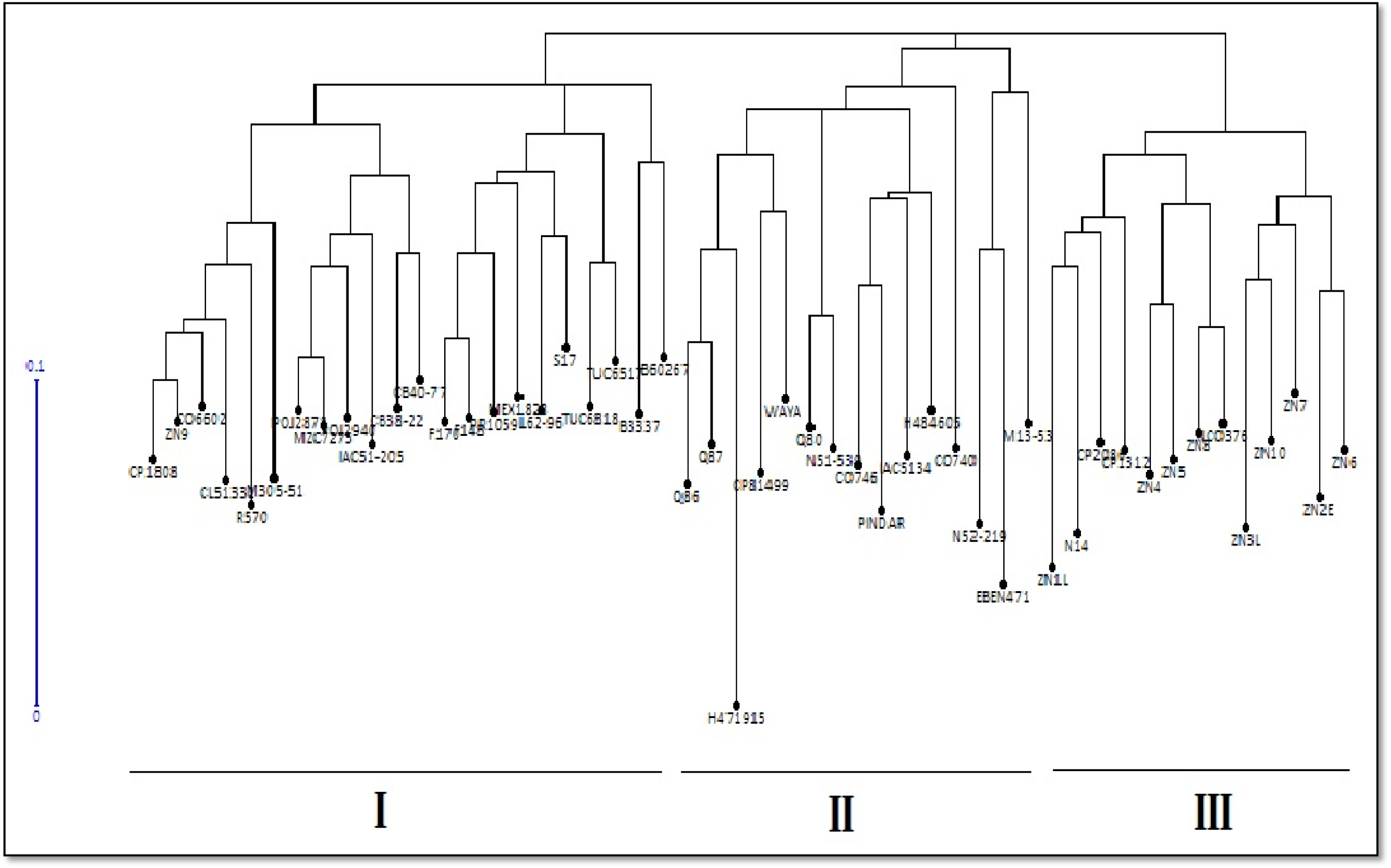
A phylogenetic tree showing clustering patterns among 50 sugarcane genotypes after SSR analyses (DARwin 6.0)

Almost all sugarcane cultivars which are being grown in the world today are derived from a few common ancestral clones from early nobilization efforts in Java (Arceneaux, 1967). India is also considered to be one of the centers of origin for *S. spontaneum* (Daniels *et al.*, 1975). This explains why the ‘POJ’ varieties were aligned together with other ancient varieties like S17 and the ‘TUC’ varieties. Indian cultivars have been introduced, adopted and utilized in various countries in the world which means that the good genomes of the ‘Co’ varieties have been integrated into the local commercial sugarcane varieties in these various countries.

From the study, Indian varieties were found in all three major clusters of the phylogenetic tree. Previous findings have found that elite sugarcane cultivars from the US, China and India shared a greater than 58% genetic similarity (Liang *et al*., 2010: Pan *et al.*, 2003a, b). Cluster III is composed of mostly genotypes of Indian and American origins. Since the environment also comes into play during variety adaptation and development, different geographical locations and breeding strategies in that same country may give rise to different chromosomal makeup of the cultivar in the same country. The long genetic distances between cluster I and III indicate different ancestry, different breeding strategies during development or incidence of mutations along the way. This information provides the breeder with a stable inference as to which genotypes to choose when making a cross so as to realize hybrid vigor and heterosis.

The consistent clustering of the Zimbabwean released varieties indicates a common selection strategy among the breeders from the countries where the varieties were imported. The Zimbabwe Sugar Industry has been importing and introducing varieties from other countries and all these varieties which have been successfully adopted into the industry were huddled in one cluster.

The three clusters were also arranged according to time of release or ‘age’ of the accessions. The clusters were arranged in descending order with the oldest accessions in cluster I which were released before 1950 (Arceneaux, 1967) and the latter (cluster II) being released pre and post 1966. For instance, the POJ varieties are early 19^th^ century varieties which were frequently used as parents in early crosses with Noble canes like Black Cheribon, Chunnee and *Saccharum officinarum* (Arceneaux, 1967). Cluster III conveniently comes last which includes varieties which were developed and released around 1995 (ZSAES Variety Guide).

In the early 19^th^ century, breeding systems had a narrow parental pool because of fewer hybrids developed compared to modern day (Milbourne *et al*., 1997). A few high performing varieties would be frequently used as parents as shown in Cluster I which was mostly comprised of POJ varieties and their offspring for instance, S17 (POJ2878 × F28) and MZC74/275 (POJ2878 × CP57-603). CB40-77 and CB40-77 are Brazilian varieties and they both have POJ2878 as a parent (ZSAES Biannual Report, 1978-1979). The POJ varieties were being crossed with Indian varieties hence presence of ‘Co’ varieties in that cluster.

## Conclusions and recommendations

This study which was carried out on only 50 of the 746 varieties present in the ZSI’s gene bank, was the first of its kind in sugarcane in Zimbabwe. Therefore, the study serves as a springboard on which further molecular and morphological characterization can be done. Greater yet unknown genetic diversity may exist amongst the rest of the varieties. Another study with a larger sample size would enable the breeder to select a mini core selection from the gene bank which would comprise of diverse varieties which would be actively useful in the breeding program and hence managed separately from the rest of the gene bank varieties. In addition to the phenotyping study, agronomic traits should be characterized so as to have a compound picture of the gene bank. Furthermore, the information gathered serves as rationale for more variety importations from international gene banks and other cane growing countries. The number of traits needs to be increased since a limited choice of traits only gives a picture of diversity in form of the traits themselves and not an overall image of disparity among the accessions. The clusters which were observed are an inference on the relationships among the potential breeding material. The breeder can utilize these groups to select distantly related individuals as parents. With the information from this study the breeder can select parents based on trait specific bias and also the apparent ability of phenotypic expression of that particular trait by the selected genotype. The information obtained from this study also serves as an excellent start for the establishment of a variety database for the Zimbabwe Sugar Industry. Morphological data can also be used for identification purposes by farmers and other stakeholders as unique descriptors and character states can be used as markers to identify or distinguish a particular variety. The study serves as a starting point for further prebreeding work to be done preliminarily before the initiation of crosses in the ZSI.

## Acknowledgements

Special acknowledgement goes to Dr. Pepukai Manjeru (Midlands State University), Dr. Leo Thokoza Mpofu (ZSAES-Chiredzi), Dr M.D.S Nzima and the ZSAES board for the financial and material support they generously and unconditionally provided. The immense contribution of Dr Washington Mutatu for laboratory, intellectual and technical assistance is greatly appreciated. Gratitude is expressed towards Mr. Manyears Chuchu and the research services field staff for the technical and intellectual expertise provided during the trials. I also acknowledge the contribution of Dr. Tinashe Muteveri and Dr Morleen Muteveri in SSR techniques training, data analysis and interpretation of results.

